# Identification of a carbohydrate-recognition motif of purinergic receptors

**DOI:** 10.1101/2023.03.06.531346

**Authors:** Lifen Zhao, Fangyu Wei, Xinheng He, Hualiang Jiang, Liuqing Wen, Xi Cheng

## Abstract

As a major class of biomolecules, carbohydrates play indispensable roles in various biological processes. However, it remains largely unknown how carbohydrates directly modulate important drug targets, such as G-protein coupled receptors (GPCRs). Here, we employed P2Y purinoceptor 14 (P2Y14), a drug target for inflammation and immune responses, to uncover the sugar nucleotide activation of GPCRs. Integrating molecular dynamics simulation with functional study, we identified the uridine diphosphate (UDP)-sugar-binding site on P2Y14, and revealed that a UDP-glucose might activate the receptor by bridging the transmembrane helices (TM) 2 and 7. Between TM2 and TM7 of P2Y14, a conserved salt bridging chain (K^2.60^-D^2.64^-K^7.35^-E^7.36^, KDKE) was identified to distinguish different UDP-sugars, including UDP-glucose, UDP-galactose, UDP-glucuronic acid and UDP-N-acetylglucosamine. We identified the KDKE chain as a conserved functional motif of sugar binding for both P2Y14 and P2Y purinoceptor 12 (P2Y12), and then designed three sugar nucleotides as agonists of P2Y12. These results not only expand our understanding for activation of purinergic receptors but also provide insights for the carbohydrate drug development for GPCRs.

## Introduction

As significant components of the organism, carbohydrates play indispensable roles in energy supply, cell signaling and immune responses (*Gagneux & Varki, 1999*). Dysregulation of carbohydrates has been proved to be associated with the development of various diseases (*Reily et al, 2019*). However, it is still elusive how carbohydrates directly act on major therapeutic targets, including G-protein coupled receptors (GPCRs) (*Cheng & Jiang, 2019; Hauser et al, 2017*). P2Y purinoceptor 14 (P2Y14) represents an outstanding model system for understanding carbohydrate-modulation of GPCRs. It belongs to P2Y purinoceptor subfamily, consisting of receptors responding to nucleotides, including adenosine diphosphate (ADP) and uridine diphosphate (UDP) (*Ralevic & Burnstock, 1998*). Distinct from the other purinoceptors, P2Y14 is potently activated by UDP and a class of carbohydrates, i.e., UDP-sugars (*Abbracchio et al, 2006; Jacobson et al, 2020*). UDP-sugars activate P2Y14 with a relative potency order of UDP-glucose (UDP-Glc), UDP-galactose (UDP-Gal), UDP-glucuronic acid (UDP-GlcA) and UDP-N-acetylglucosamine (UDP-GlcNAc) (*Chambers et al, 2000; Hamel et al, 2011; Ko et al, 2009; Ko et al, 2007*). These sugar nucleotides act as important signaling molecules via P2Y14 to mediate many physiological processes (*Amison et al, 2017; Breton & Brown, 2018; Ferreira et al, 2017; Lazarowski, 2010; Muller et al, 2005; Sesma et al, 2016*). Particularly, UDP-Glc regulates immune responses and associate with asthma, kidney injury, and lung inflammation (*Amison et al., 2017; Breton & Brown, 2018; Ferreira et al., 2017; Muller et al., 2005; Sesma et al., 2016*). As an isomer of UDP-Glc, UDP-Gal is present in various cell models, including physiologically relevant primary cultures of human bronchial epithelial cells (*Lazarowski, 2010*). It remains unknown how these sugar nucleotides are recognized by P2Y14.

As the closest homolog to P2Y14, P2Y purinoceptor 12 (P2Y12) has not been reported to be activated by any sugar nucleotide (*Jacobson et al., 2020; Ralevic & Burnstock, 1998*). P2Y12 is potently activated ADP. The reported agonist-bound structures of P2Y purinoceptor 12 (P2Y12) provide insights to understand the nucleotide activation of P2Y purinoceptors. The crystal structures of P2Y12 show that a full agonist 2-methylthio-adenosine-59-diphosphate (2MeSADP, a close analogue of ADP) binds to an extracellular pocket consisting of transmembrane (TM) helices (*Zhang et al, 2014a*). Since P2Y12 is highly similar to P2Y14 with 45.67% amino acid sequence identity, it would be interesting to investigate whether this receptor is also sensible to sugar nucleotides.

Here, we combined molecular docking, molecular dynamics (MD) simulations and functional study to reveal the molecular mechanism how P2Y14 is activated by a sugar nucleotide. The ligand-binding models of different UDP-sugars (UDP-Glc, UDP-Gal, UDP-GlcA and UDP-GlcNAc) were quantitatively characterized to identify the sugar-recognition site of P2Y14. Both P2Y14 and P2Y12 were employed to unveil a conserved sugar-binding motif. Multiple carbohydrates were designed and validated as their agonists targeting the conserved functional motif.

## Results

### Identification of sugar-binding site in P2Y14

Both UDP and UDP-Glc potently activate P2Y14 with EC50 values of 50.9 ± 6.1 nM and 40.3 ± 1.5 nM, respectively (***Figure 1 A-C***). Compared with UDP, UDP-Glc showed an increased potency on P2Y14 at high concentration (***Figure 1B-C***), suggesting that the sugar moiety of UDP-Glc contributes to activating P2Y14. To investigate how UDP-Glc regulates the P2Y14 via its sugar moiety, we used molecular docking to construct UDP-Glc-bound models of P2Y14 and compared them with UDP-bound P2Y14 models (***Figure 1D-G***). Because the protein structure of P2Y14 is unrevealed, we employed the X-ray structures of P2Y12 (*Zhang et al., 2014a*) as templates to constructed homology models of human P2Y14. The molecular docking showed both UDP and UDP-Glc bind to an extracellular pocket consisting of transmembrane (TM) helices 2-7 and extracellular loop (ECL) 2 (***Figure 1D***), which is corresponding to a known agonist binding pocket of P2Y12 (*Zhang et al., 2014a*). The docking score of UDP-Glc is −9.3 kcal/mol, which is 0.8 kcal/mol lower than that of UDP (docking score = −8.5 kcal/mol), indicating that both UDP and UDP-Glc stably bind to P2Y14.

**Figure 1.**
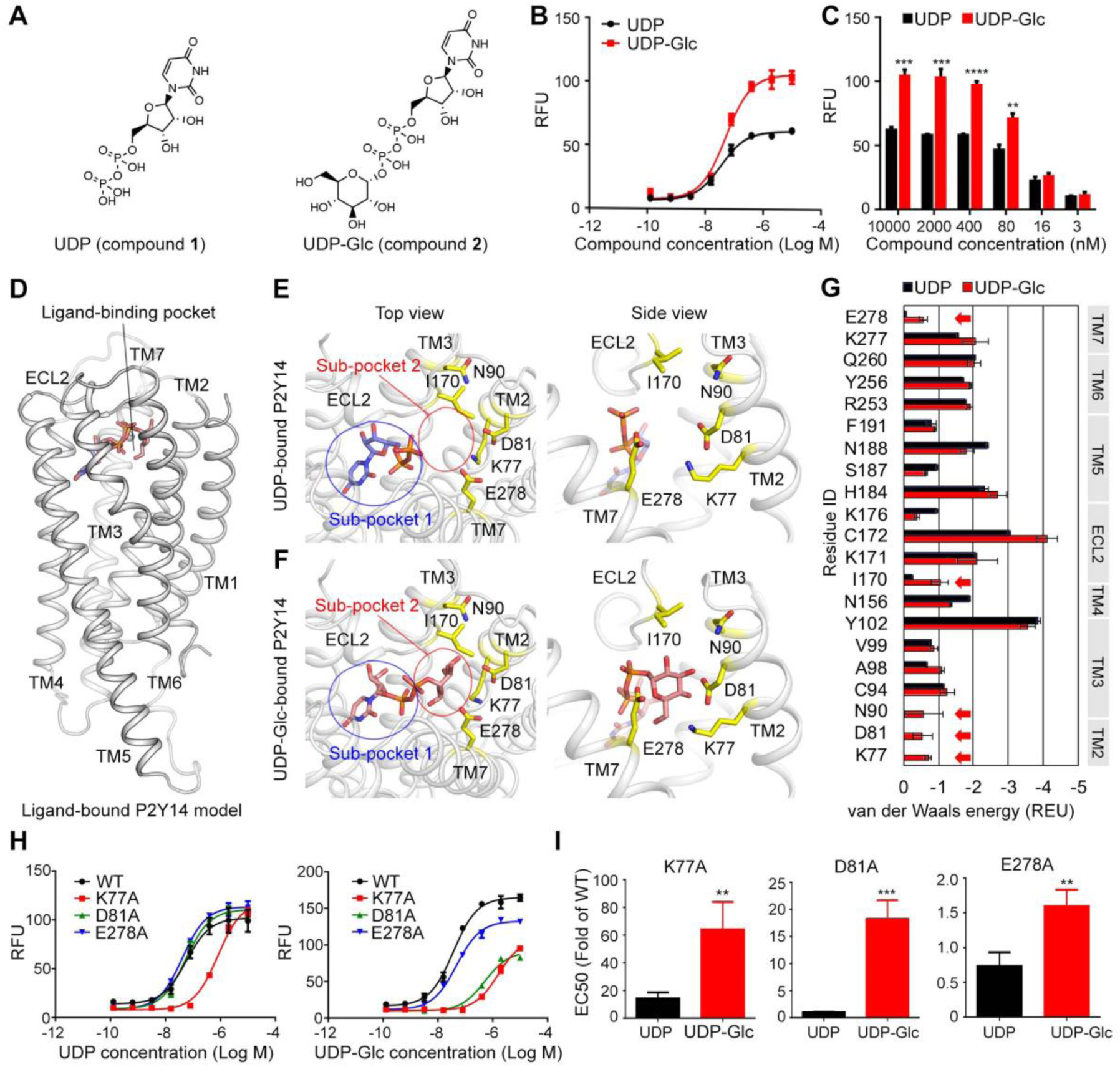
Identification of sugar-binding sites in P2Y14. (**A**) Chemical structures of UDP and UDP-Glc. (**B**) Concentration-response curves of calcium mobilization for UDP or UDP-Glc in HEK293 cells transiently co-transfected with human P2Y14 and Gα_qi5_. Data are shown as mean ± SEM (*n* = 3). See also Figure 1***—source data 1*** and ***Supplementary file 1***. (**C**) Concentration-dependent comparation of calcium mobilization for UDP and UDP-Glc in HEK293 cells transiently co-transfected with human P2Y14 and Gα_qi5_ (*n* = 3); \*\**P* < 0.01, \*\*\**P* < 0.001, \*\*\*\**P* < 0.0001. (**D**) Ligand-bound model of P2Y14. Protein and compound are shown in cartoon and stick representations. (**E, F**) Docking models of UDP (**E**) and UDP-Glc (**F**) to P2Y14. Key residues are highlighted in yellow. Two sub-pockets for ligand binding are marked with circles. (**G**) Decomposition of ligand-binding energy for each receptor residue (*n* = 10). (**H**) Calcium mobilization concentration-response curves for UDP or UDP-Glc in HEK293 expressing P2Y14 WT and mutants (*n* = 3). See also Figure 1***—source data 2*** and ***Supplementary file 1***. (**I**) Comparation of EC50s for UDP-Glc and UDP in HEK293 cells expressing P2Y14 mutants in calcium mobilization assay (*n* = 3); \*\**P* < 0.01, \*\*\**P* < 0.001. See also Figure 1***—source data 3***. **Source data 1.** Potency of UDP or UDP-Glc in HEK293 cells expressing P2Y14. **Source data 2.** Potency of UDP or UDP-Glc in HEK293 cells expressing P2Y14 WT and mutants. **Source data 3.** Comparation of EC50s for UDP-Glc and UDP in HEK293 cells expressing P2Y14 mutants.

Compared with the UDP-bound receptor model (***Figure 1E***), the UDP-Glc-bound model showed extra interactions between the glucose-moiety and the TM2, TM3, TM7 and ECL2 of P2Y14 (***Figure 1F***), enhancing the binding of UDP-Glc. Based on these molecular docking models, we further decomposed the ligand-binding energy to each receptor residue (***Figure 1G***). Five residues (K77^2.60^, D81^2.64^, N90^3.21^, I170^ECL2^ and E278^7.36^; superscript indicates Ballesteros-Weinstein residue numbering (*Ballesteros, 1995*)) were predicted to stabilize UDP-Glc binding (***Figure 1F, G***), while they made few energetic contributions (van der Waals energy > −0.25 Rosetta energy unit) to UDP binding (***Figure 1E, G***). As shown in Fig 1E, F, two sub-pockets of P2Y14 were unveiled for ligand binding. The sub-pocket 1 is formed by 16 residues of TMs 3-7 and ECL2 (***Figure 1G, Figure 1—Figure supplement 1A, B***) and binds to the nucleotide moiety of the agonist, i.e., UDP. The sub-pocket 2 is the specific sugar-binding site involving K77^2.60^, D81^2.64^, N90^3.21^, I170^ECL2^ and E278^7.36^ (***Figure 1F, G***). These residues are primarily charged or polar amino acids, which could made hydrogen bonds with the glucose hydroxyl groups of UDP-Glc (***Figure 1F***). To validate the proposed sugar-binding sites, we designed single-point mutations of these five residues (K77A, D81A, N90A, I170A and E278A). Among these mutants, D81A and E278A showed significantly reduced activities by UDP-Glc compared with the wild-type (WT) group (***Figure 1H***). However, substitution of D81^2.64^ or E278^7.36^ by alanine did not significantly affect the receptor activities by UDP (***Figure 1H***). Interestingly, K77A mutation diminished both UDP-Glc- and UDP-induced calcium mobilization (***Figure 1H***), but it showed greater impact on UDP-Glc-induced receptor responses than UDP-induced ones (***Figure 1H, I***), suggesting extra interactions between K77^2.60^ and sugar-moiety of UDP-Glc. Two mutations on TM3 and ECL2 (N90A and I170A) did not significantly affect the receptor responses by UDP or UDP-Glc (***Figure 1—Figure supplement 1C, D***). These findings indicate that sub-pocket 2 residues of TM2 and TM7 provide major contributes to stabilizing the sugar moiety of UDP-Glc.

### UDP-Glc as a “glue” for P2Y14 activation

The molecular docking employs rigid side chains of the receptor and does not include the influence of explicit environment on molecular interactions. To investigate how UDP-Glc interacts with P2Y14, we performed all-atom MD simulations of the P2Y14 receptor with and without UDP-Glc (***Figure 2A***). We used the molecular docking model of P2Y14 to construct the simulation systems. Apo P2Y14 and UDP-Glc-bound P2Y14 simulation models showed different conformations in TM6 and TM7 (***Figure 2A, B***). In UDP-Glc-bound P2Y14 simulations, the extracellular tip of TM6 shifted over 3 Å and TM7 over 4 Å towards the receptor core, compared with the apo P2Y14 simulations (***Figure 2A, B***). This inward shift of TM6 and TM7 allowed formation of polar and ionic interactions with the UDP-Glc (***Figure 2B, C***). During UDP-Glc-bound P2Y14 simulations, two charged residues K277^7.35^ and E278^7.36^ formed hydrogen bonds with the glucose 6’ hydroxyl group of UDP-Glc to keep TM7 close to the receptor core (***Figure 2B, C***), and an arginine residue (R253^6.55^) formed a salt bridge with the phosphate group of UDP-Glc to stabilize the inward shift of TM6 (***Figure 2B, C***). Consistently, compared with WT group (EC50 of 40.3 ± 1.5 nM), single-point mutations (R253A and E278) of TM6 and TM7 helices resulted to diminished UDP-Glc-induced calcium mobilization (EC50 of 808.6 ± 43.6 nM for R253A and 60.2 ± 3.6 nM for E278) (***Figure 1H, Figure 2D***). In addition, at the extracellular side, the distance between TM5 and TM6 of UDP-Glc-bound P2Y14 was 5.9 Å shorter than that in the apo system (***Figure 2—Figure supplement 1A, B***). Y189^5.41^ and T257^6.59^ made stable hydrophobic interactions to maintain the tight compact between TM5 and TM6 in UDP-Glc-bound-P2Y14 simulations, while TM6 did not interact with TM5 at the extracellular side in the apo simulations (***Figure 2—Figure supplement 1A, B***). Compared with WT group (EC50 of 40.3 ± 1.5 nM), a mutation of TM6 (T257A) showed significantly reduced UDP-Glc-induced responses (EC50 of 504.9±15.9 nM) (***Figure 2—Figure supplement 1C***), fully agreeing with our simulation models. Collectively, these data suggest UDP-Glc might serve as intramolecular “glue” to make a tight helical bundle of P2Y14, involving TM6 and TM7.

**Figure 2.**
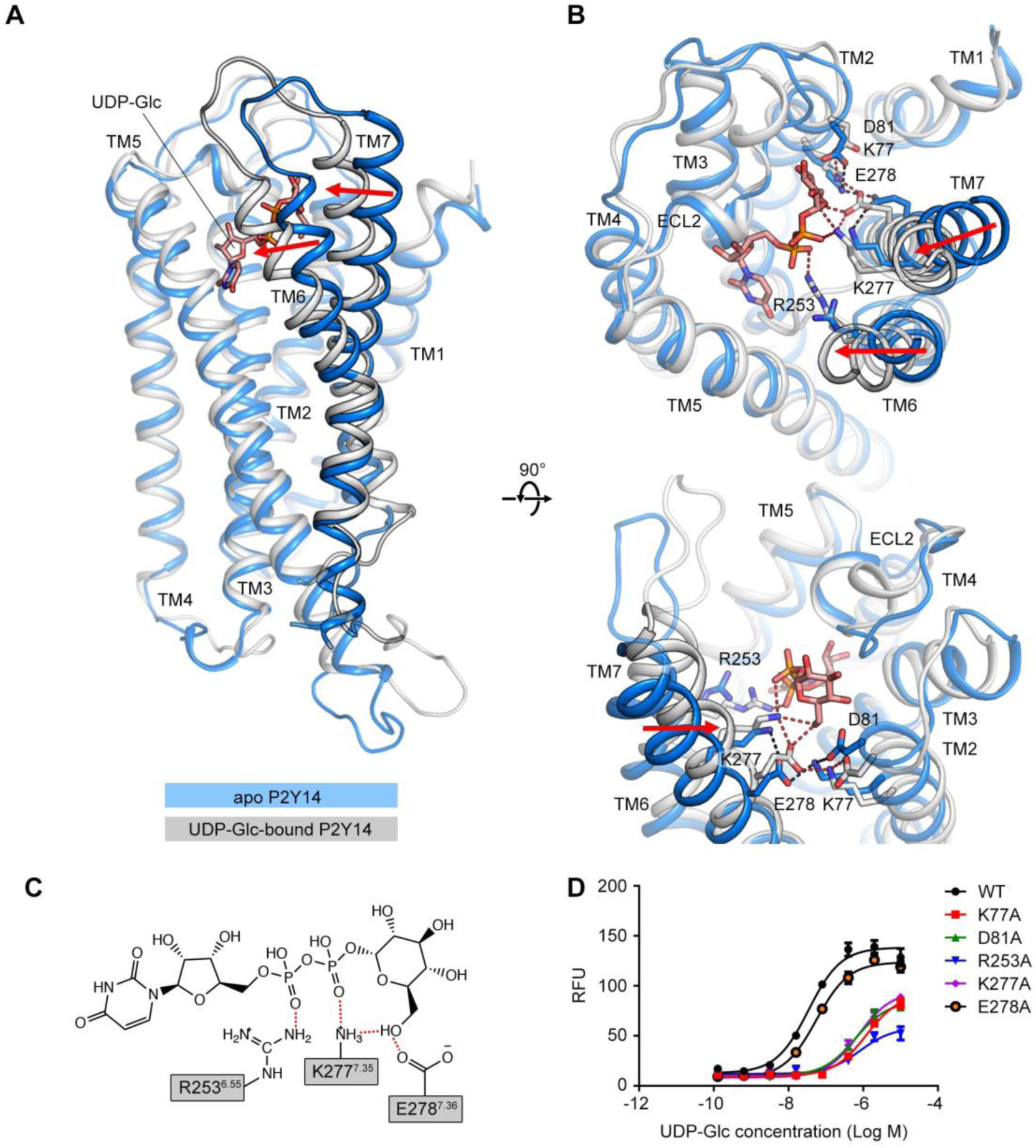
Comparison of the apo P2Y14 and UDP-Glc-bound P2Y14 simulation systems. (**A-B**) Side (**A**) and top (**B**) views of representative models of apo P2Y14 and UDP-Glc-bound P2Y14. Receptor is shown as cartoon. Ligand and key residues are shown as sticks. Movement of the extracellular tips of TM6 and TM7 towards the receptor core is shown by arrows. See ***Supplementary file 2*** for computational characterization of conformational changes. (**C**) Key interactions between P2Y14 and UDP-Glc. Hydrogen bonds and salt bridges are displayed as red dashed lines. See ***Supplementary file 2*** for pairwise interaction details. (**D**) Concentration-response curves of calcium mobilization for UDP-Glc in HEK293 expressing P2Y14 WT and mutants. Data are shown as mean ± SEM (*n* = 3). See also Figure 2***—source data 1*** and ***Supplementary file 1***. **Source data 1.** Potency of UDP-Glc in HEK293 expressing P2Y14 WT and mutants.

### Molecular recognition of P2Y14 via sugar-binding site

P2Y14 could be activate by different UDP-sugars with distinct potencies. With only one group substitution at the sugar moiety, UDP-Glc induced stronger activity on P2Y14 (EC50 = 40.3 ± 1.5 nM) than the other UDP-sugars, i.e., UDP-Gal (EC50 = 78.3 ± 9.2 nM), UDP-GlcA (EC50 = 59.9 ± 4.8 nM) and UDP-GlcNAc (EC50 = 184.4 ± 11.8 nM) (***Figure 3A, B***. To investigate how P2Y14 recognizes different sugar moieties, we performed MD simulations of the human P2Y14 receptor complex with UDP-Gal, UDP-GlcA and UDP-GlcNAc, respectively, and compared them with the UDP-Glc-bound P2Y14 simulations. We observed that UDP-Gal, UDP-GlcA and UDP-GlcNAc bound to P2Y14 at the same pocket as UDP-Glc. Similar to UDP-Glc, their uridine groups occupied the sub-pocket 1 of P2Y14, while their sugar moieties bound to the sub-pocket 2 during simulations (***Figure 3C-F***). At the sub-pocket 2, a stable salt bridging chain formed by four charged residues (K77^2.60^, D81^2.64^, K277^7.35^ and E278^7.36^) were observed in all systems (***Figure 3C-F***). The negative charged glutamic acid residue E278^7.36^ linked TM2 and TM7 helices by forming salt bridges with K77^2.60^ and K277^7.35^, while the other negative charged residue D81^2.64^ forming a salt bridge with K77^2.60^ to further stabilize these ionic interactions (***Figure 3C-F***).

**Figure 3.**
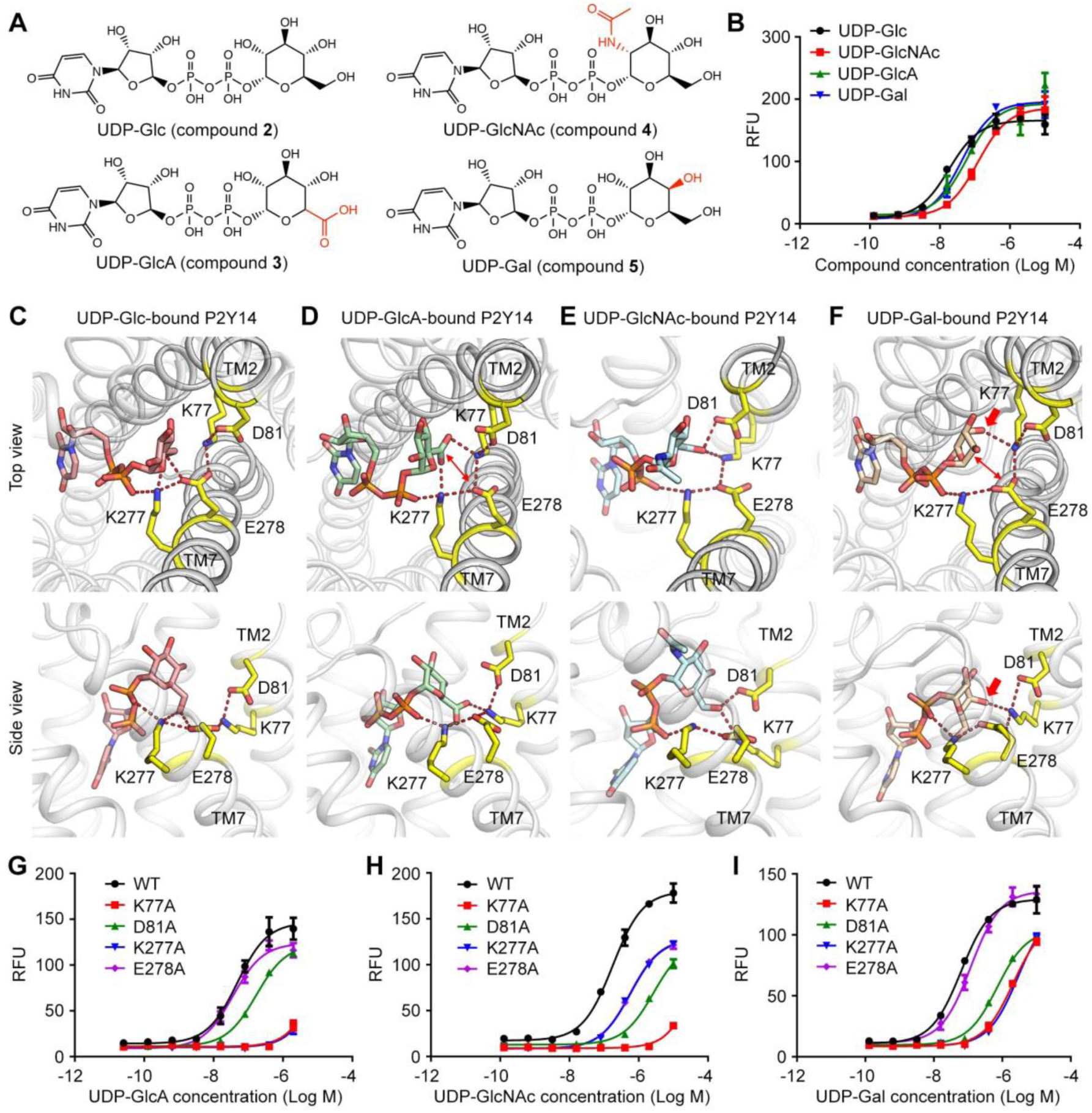
Sugar moiety recognition of P2Y14. (**A**) Chemical structures of UDP-Glc, UDP-GlcA, UDP-GlcNAc and UDP-Gal. (**B**) Concentration-response curves of calcium mobilization for different UDP-sugars in HEK293 cells transiently co-transfected with human P2Y14 and Gα_qi5_. Data are shown as mean ± SEM (*n* = 3). See also Figure 3***—source data 1***. (**C-F**) Molecular recognition of P2Y14 for UDP-Glc (**C**), UDP-GlcA (**D**), UDP-GlcNAc (**E**) and UDP-Gal (**F**). Receptor, ligands and key residues are shown in cartoon and stick representations. Hydrogen bonds and salt bridges are displayed as red dashed lines. See ***Supplementary file 2*** for pairwise interaction details. (**G-I**) Concentration-response curves of calcium mobilization for UDP-GlcA (**G**), UDP-GlcNAc (**H**) and UDP-Gal (**I**) in HEK293 expressing P2Y14 WT and mutants. Data are shown as mean ± SEM (*n* = 3). See also Figure 3***—source data 2***. **Source data 1.** Potency of UDP-GlcA, UDP-GlcNAc and UDP-Gal in HEK293 expressing P2Y14. **Source data 2.** Potency of UDP-GlcA, UDP-GlcNAc and UDP-Gal in HEK293 expressing P2Y14 WT and mutants.

In simulations, different sugar moieties bound to the K77^2.60^-D81^2.64^-K277^7.35^-E278^7.36^ salt bridging chain with distinct binding modes (***Figure 3C-F, Figure 3—Figure supplement 1***). For UDP-Glc, both K277^7.35^ and E278^7.36^ could form hydrogen bonds with the glucose 6’ hydroxyl group to keep TM7 close to the receptor core (***Figure 3C, Figure 3—Figure supplement 2A***). However, in UDP-GlcA-bound P2Y14 simulations, at the corresponding position, the 5’ carboxyl group of the sugar moiety repelled the negatively charged E278^7.36^ (***Figure. 3D, Figure 3—Figure supplement 2B***). Compared with that of WT group (EC50 = 59.9 ± 4.8 nM), the single-point mutation of E278A significantly enhanced the UDP-GlcA-induced calcium mobilization with a EC50 of 38.2 ± 2.2 nM (***Figure 3G***). These experimental results support with the proposed sugar-binding model (***Figure 3D***) and suggest that the reduced interactions of UDP-GlcA with TM7 (E278^7.36^) might contributed to its weak potency on P2Y14. Substitution of 2’ hydroxyl group by an acetamido group led to a rotation of the sugar moiety of UDP-GlcNAc in simulations (***Figure 3E***). Consequentially, the 6’ hydroxyl of N-acetylglucosamine group flipped to form hydrogen bonds with K77^2.60^ and D81^2.64^ instead of K277^7.35^ and E278^7.36^ (***Figure 3E, Figure 3—Figure supplement 2C***). Consistently, single-point mutation of D81A made more significant effect to reduce the UDP-GlcNAc-induced receptor activities, compared with that of E278A (***Figure 3H***). Compared with the other three UDP-sugars, UDP-Gal has a different orientation of 4’ hydroxyl group. The 4’ hydroxyl group of galactose formed a stable hydrogen bond with K77^2.60^ and disrupted the interaction between 6’ hydroxyl group with E278^7.36^ (***Figure 3F, Figure 3—Figure supplement 2D***). Compared with UDP-Glc, UDP-Gal had more interactions with TM2 and less interactions with TM7 (E278^7.36^). Substitute of E278^7.36^ by alanine did not significantly affect the UDP-Gal-induced receptor response (***Figure 3I***), agreeing with the proposed UDP-Gal-binding model (***Figure 3F***). For all UDP-sugars, at least three residues of K77^2.60^, D81^2.64^, K277^7.35^ and E278^7.36^ participated in ligand binding (***Figure 2, Figure 3***). Both computational models and experimental data indicate the K77^2.60^-D81^2.64^-K277^7.35^-E278^7.36^ salt bridging chain as a sugar-binding site of P2Y14, which can recognize different sugar moieties. The interactions of ligands with the TM7 might determine the ligand potency on P2Y14.

### Conserved sugar-binding motif for P2Y12 and P2Y14

In previous sections, we have identified K77^2.60^-D81^2.64^-K277^7.35^-E278^7.36^ salt bridging chain as an important functional site for sugar moiety recognition and UDP-sugar activation of P2Y14. These four residues (K^2.60^, D^2.64^, K^7.35^ and E^7.36^) are conserved between P2Y14 and its closest homolog, i.e., P2Y12 (***Figure 4A, Figure 4—supplement 1A***). Therefore, we assumed P2Y12 also can be activated by carbohydrate ligands. P2Y12 is activated by ADP (*Herbert & Savi, 2003*), but it has not been reported to be activated by any carbohydrate. To validate our assumption, we designed and synthesized three ADP-sugars, i.e., ADP-glucose (ADP-Glc), ADP-glucuronic acid (ADP-GlcA) and ADP-mannose (ADP-Man), and then tested whether they can activate P2Y12 (***Figure 4B, C***). We docked ADP-Glc and ADP-Man to the X-ray structure of P2Y12 (*Zhang et al., 2014a*). The docking scores are −9.4 kcal/mol for ADP-Glc, for −10.0 kcal/mol for ADP-GlcA and −9.3 kcal/mol for ADP-Man (***Figure 4—supplement 1B***), suggesting they stably bound to P2Y12. Consistently, in calcium mobilization assays, ADP-Glc, ADP-GlcA and ADP-Man activated P2Y12 with EC50 values of 3.4 ± 0.4 μM, 1.3 ± 0.1 μM and 12.3 ± 0.9 μM, respectively (***Figure 4C***). Single-point mutations of K80^2.60^, D84^2.64^, K280^7.35^ and E281^7.36^ significantly diminished ADP-Glc-, ADP-GlcA- and ADP-Man-induced responses, compared with WT P2Y12 (***Figure 4C***). These findings not only validate our assumption that P2Y12 can be activated by sugar nucleotides, but also indicate the conserved K^2.60^-D^2.64^-K^7.35^-E^7.36^ (KDKE) salt bridging chain as a functional motif for sugar binding.

**Figure 4.**
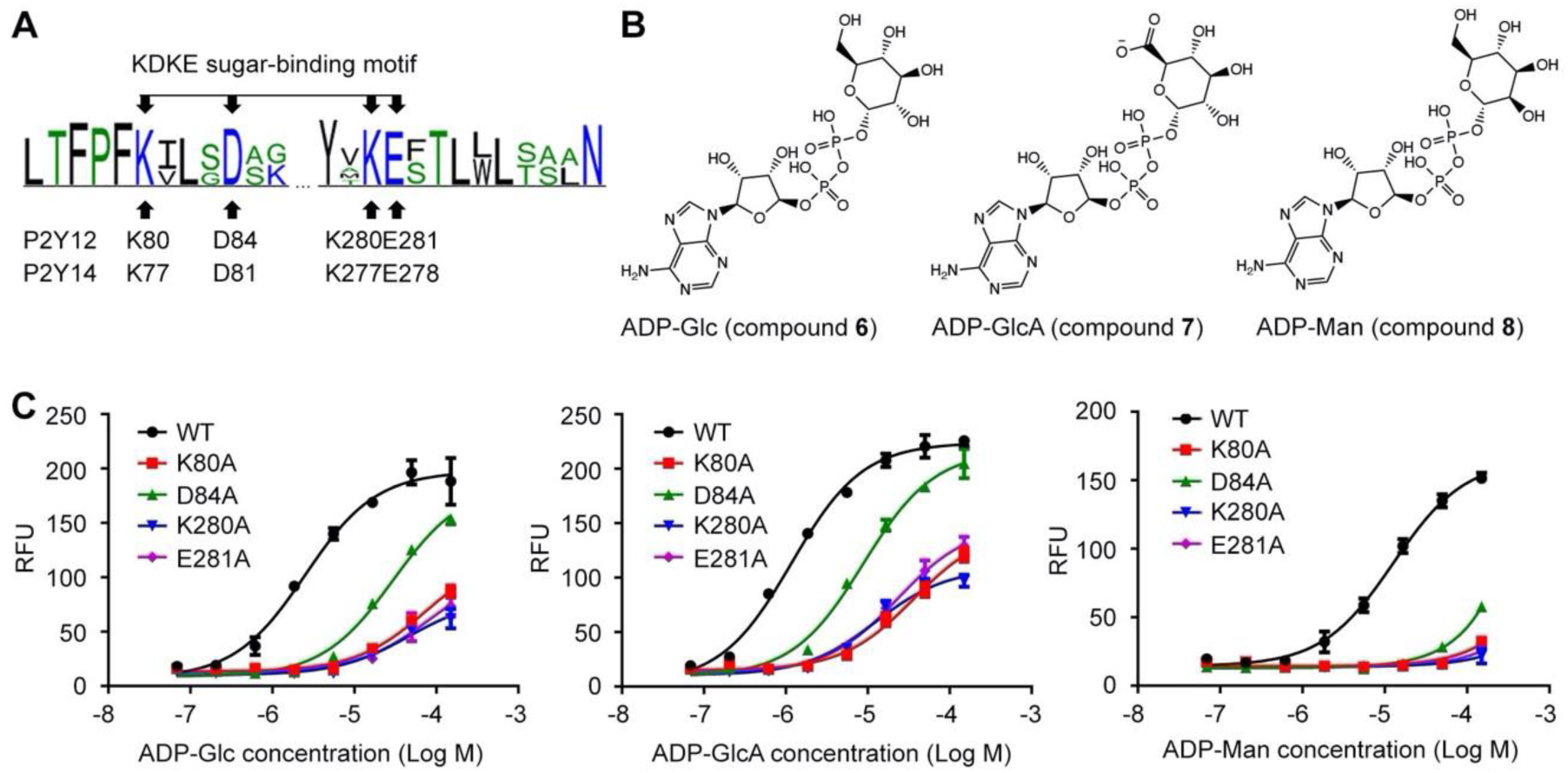
ADP-sugars binding to P2Y12. (**A**) Sequence log of the alignment between P2Y12 and P2Y14. Sequences of P2Y12 and P2Y14 involving 379 species were collected for making sequence alignments. See ***Supplementary file 3*** for species repertoire information. The height of a letter is proportional to the relative frequency of that residue at a particular site. Four residues of KDKE sugar-binding motif are marked by arrows, with the corresponding residues in P2Y12 and P2Y14. (**B**) Chemical structure of ADP-Glc, ADP-GlcA and ADP-Man. (**C**) Calcium mobilization concentration-response curves for ADP-Glc, ADP-GlcA and ADP-Man in HEK293 expressing P2Y12 WT and mutants. Data are shown as mean ± SEM (*n* = 3). See also Figure 4***—source data 1*** and ***Supplementary file 1***. **Source data 1.** Potency of ADP-Glc, ADP-GlcA and ADP-Man in HEK293 expressing P2Y12 WT and mutants.

## Discussion

Mediated by UDP-sugars, P2Y14 plays an important role in immune responses and inflammation (Arase et al, 2009; Barrett et al, 2013; Breton & Brown, 2018; Ferreira et al., 2017; Muller et al., 2005; Sesma et al, 2012; Sesma et al., 2016), and possibly insulin resistance (Wang et al, 2008). Breton et al. found that kidney collecting duct intercalated cells present high levels of P2Y14, which is activated by UDP-Glc to promote neutrophil infiltration and renal inflammation (Breton & Brown, 2018). Exerting excessive P2Y14-mediated inflammatory reactions, high concentration of UDP-sugars was observed in extracellular tissue surrounding airway epithelial cells and lung secretions of cystic fibrosis patients (Ferreira et al., 2017; Muller et al., 2005; Sesma et al., 2016). UDP-Glc is also released from liver cells in obese states, possibly via hepatocellular apoptosis, leading to liver inflammation and insulin resistance (Wang et al., 2008). All these results indicate importance of UDP-sugar regulation of P2Y14 in pathological progresses.

In this work, we built the molecular model of UDP-Glc-bound P2Y14 to answer the long-standing question of sugar nucleotide-regulation of the purinergic receptor. Binding to an extracellular pocket involving TMs 2 and 7 (***Figure 1***), the UDP-Glc might serve as intramolecular “glue” attaching to TM6 and TM7 to activate P2Y14 (***Figure 2***). The agonist-induced remarkable conformational changes of TM6 and TM7 are also reported for P2Y12 (*Zhang et al., 2014a*). Compared with the AZD1283-bound (antagonist-bound) P2Y12 structure (*Zhang et al, 2014b*), the extracellular part of TM6 in the 2MeSADP-bound (agonist-bound) P2Y12 structure shifts over 10 Å and TM7 over 5 Å towards the center of TM helix bundle (*Zhang et al., 2014a*). The close parallel of P2Y12 and P2Y14 in the agonist-induced conformational changes indicates a common ligand-induced activation mechanism shared by purinergic receptors. In addition, in the studies involving the other UDP-sugars, we also found that the interactions between sugar-moieties of agonists with TM7 (E278^7.36^) is determinant for UDP-sugars’ potencies (***Figure 3***).

The carbohydrate-binding site has not been fully characterized for GPCRs. Except for P2Y14, it has not been reported that the other members of P2Y12-like subfamily can be directly activated by carbohydrates. Integrated computational modeling with mutagenesis study, we identified a conserved carbohydrate-binding motif (KDKE) for both P2Y14 and P2Y12 (***Figure 4***). The KDKE motif not only participates in receptor activation by bridging TM2 and TM7 (K77^2.60^, D81^2.64^, K277^7.35^ and E278^7.36^) (***Figure 2***), but also recognize different sugar moieties, including glucose, galactose, glucuronic acid and N-acetylglucosamine groups (***Figure 3, Figure 4***). Remarkably, this KDKE motif can distinguish isomers as UDP-Glc and UDP-Gal. Our MD simulations showed that KDKE motif attracted the 6’ hydroxyl group of glucose but the 4’ hydroxyl group of galactose (***Figure 3C, F***). Consistent with our observations, a previous structure-activity relationship study revealed that selective mono-fluorination of the 6’ hydroxyl group of the glucose moiety results to 4-fold less potency on P2Y14 (*Ko et al., 2009*). As another member of P2Y12-like subfamily, P2Y13 also has the conserved K^2.60^-D^2.64^-K^7.35^-E^7.36^ site (***Figure 4—supplement 2***), suggesting it might be regulated by carbohydrates. GPR87 is a close homolog of P2Y14 with the sequence identity of 44.94%. GPR87 has K/R^2.60^, D^2.64^, K/E^7.35^ and E^7.36^ at the corresponding positions of the KDKE sugar-binding motif, indicating varied carbohydrate-sensitivities of this receptor in different species (***Figure 4—supplement 3***). Similar to the KDKE motif of the receptors, the UDP-sugar-binding sites consisting of charged residues have be discovered for sugar transferases (*Gerlach et al, 2018; Hao et al, 2021*). In typical glycosylation transfers as TarP and SseK3, two aspartic acids and one positively charged residue (arginine or lysine) participate in recognition of 3’ or 4’ hydroxyl groups of GlcNAc or GalNAc moiety (*Gerlach et al., 2018; Hao et al., 2021*). However, a salt bridging chain has not been observed in these sugar-binding sites. The different arrangements of UDP-sugar-binding sites between P2Y14 and these sugar transferases might be determinant to their sugar selectivity.

In conclusion, we revealed a conserved carbohydrate binding motif in both P2Y12 and P2Y14, extending our understanding how carbohydrates regulate GPCRs. Our molecular models of different sugar nucleotides provide great details for carbohydrate-activation and recognition of these receptors, which would inspire further carbohydrate drug development for GPCRs. Whether the other carbohydrate-binding motifs exist in GPCRs is currently unknown. Further investigations focused on carbohydrate-regulation of GPCRs will continue to add both new concepts and physiological understanding to the field.

## Materials and Methods

### Chemicals

UDP-GlcNAc was prepared from D-GlcNAc as reported previously (*Zheng et al, 2022*). UDP-Glc and UDP-GlcA were prepared from Sucrose (*Wang et al, 2022*). UDP-Gal was prepared from D-Gal (*Muthana et al, 2012*). ADP-Man was synthesized by a two-step strategy. In detail, Man-1-p was firstly synthesized from D-Man using NahK from Bifidobacterium longum (*Nishimoto & Kitaoka, 2007*) and ATP as phosphorylation donor. Man-1-p was purified from the reaction mixture by the silver nitrate precipitation method (*Liuqing et al, 2016; Wen et al, 2015*). Then, ADP-Man was synthesized from Man-1-p and ATP by a GDP-mannose pyrophosphorylase from Pyrococcus furiosus, which could take ATP as substrate.

### Cell culture and transient transfections

HEK293 cells were cultured in Dulbecco’s modified Eagle’s medium (DMEM) with 10% fetal bovine serum (FBS). All cells were maintained at 37℃ in humidified incubators with 5% CO_2_ and 95% air. Human P2Y14 or P2Y12 receptors and G protein α-subunit (Gα_qi5_) were transiently co-transfected into HEK293 cells using PolyJet In Vitro DNA Transfection Reagent (SignaGen) according to manufacturer’s instructions. Thus, a mixture of 1 μg of receptor DNA and 1 μg of Gα_qi5_ DNA was used to transfected into the 6-well plate cells at 90% confluency. Transiently transfected HEK293 cells were subsequently in Ca^2+^ assays at 48-hour post-transfection.

### Cell surface expression

Human P2Y14 or P2Y12 was cloned into a pcDNA3 vector with HA tag for expression in HEK293 cells. Mutants of P2Y14 or P2Y12 were constructed according to Fast Mutagenesis System (TransGen). Cell surface expression of P2Y14 or P2Y12 was analyzed by flow cytometry. HEK293 cells were transfected with pCDNA3-HA-P2Y14 or P2Y12 in 6-well plate overnight. After having been incubated with rabbit anti-HA primary antibody (1:800, CST) for 1 hour at 4 ℃, the cells were incubated with goat anti-rabbit IgG(H+L) FITC conjugate secondary antibody (1:200, TransGen) for 50 minutes at 4 ℃. Data were collected with a flow cytometer (FACS Calibur, BD) and analyzed with FlowJo software.

### Intracellular Ca^2+^ mobilization

Intracellular Ca^2+^ assays were carried out as follows. HEK293 cells were seeded (80000 cells/well) into Matrigel-coated 96-well plate 24 hour prior to assay. The cells were incubated with 2 μM Fluo-4 AM (Invitrogen) diluted in HBSS solution (0.4 gL^−1^ KCl, 0.12 gL^−1^ Na_2_HPO_4_·12H_2_O,0.06 g L^−1^ KH_2_PO_4_, 0.35 gL^−1^ NaHCO_3_, 0.14 gL^−1^ CaCl_2_, 0.10 g L^−1^ MgCl_2_·6H_2_O, 0.05 g L^−1^ MgSO_4_, and 8.0 g L^−1^ NaCl) at 37 °C for 50 minutes. After dye loading, the cells were treated with the compounds of interest. Then, calcium response (relative fluorescence unit, RFU) was measured using Flexstation 3 (Molecular Device) with fluorescence excition made at 485 nm and emission at 525 nm.

### Molecular modeling, docking and energy decomposition

Using the crystal structures of agonist-bound P2Y12 (PDB codes 4PXZ, 4PY0) (*Zhang et al., 2014a*) as templates, we employed Modeller (*Sali & Blundell, 1993*) to construct the human P2Y14 models. The human P2Y12 models are also built using these P2Y12 crystal structures (PDB codes 4PXZ, 4PY0) (*Zhang et al., 2014a*). The models with the lowest root mean square deviations from their template structures were selected for further analysis. A ligand was docked to the receptor using Schodinger Glide software in SP mode with default parameters (*Friesner et al, 2004*). A pocket binding to the ligand with Glide G-scores below −6.5 kcal/mol were considered as a possible ligand-binding pocket. To involve receptor flexibility, we used RosettaLigand (*Davis & Baker, 2009*) to generate representative ligand-bound receptor models. After Rosetta-based docking, the top 1,000 models with lowest binding energy score were selected. Then, they were further scored with the ligand-binding energy between ligand and receptor. The top 10 models with the lowest ligand-binding energy were selected for energy decomposition. The van der Waals energy of ligand binding was mapped to each receptor residue by residue_energy_breakdown utility (*Davis & Baker, 2009*). The model with the lowest ligand binding energy was used as the representative model.

### Modeling and simulations

To build a simulation system, we place the molecular model into a 1-palmitoyl-2-oleoyl-sn-glycero-3-phosphocholine lipid bilayer. The lipid embedded complex model was solvated in periodic boundary condition box (80 Å x 80 Å x 120 Å) filled with TIP3P water molecules and 0.15 M KCl using CHARMM-GUI (*Wu et al, 2014*). Each system was replicated to performed three independent simulations. On the basis of the CHARMM36m all-atom force field (*Guvench et al, 2011; Huang et al, 2017; MacKerell et al, 1998*), molecular dynamics simulations were conducted using GROMAS 5.1.4 (*Hess et al, 2008; Van Der Spoel et al, 2005*). After 100 ns equilibration, a 500-ns production run was carried out for each simulation. All productions were carried out in the NPT ensemble at temperature of 303.15 K and a pressure of 1 atm. Temperature and pressure were controlled using the velocity-rescale thermostat (*Bussi et al, 2007*) and the Parrinello-Rahman barostat with isotropic coupling (*Aoki & Yonezawa, 1992*), respectively. Equations of motion were integrated with a 2 fs time step, the LINCS algorithm was used to constrain bond length (*Hess, 2008*). Non-bonded pair lists were generated every 10 steps using distance cut-off of 1.4 nm. A cut-off of 1.2 nm was used for Lennard-Jones (excluding scales 1-4) interactions, which were smoothly switched off between 1 and 1.2 nm. Electrostatic interactions were computed using particle-mesh-Ewald algorithm with a real-space cutoff of 1.2 nm. The last 200 ns trajectory of each simulation was used to calculate average values.

### Sequence analysis

To analyze the conservation of residual sites, we collected sequences of receptors from UniProt database involving 379 species. See ***Supplementary file 3*** for species repertoire information. The multiple sequence alignments were performed using Clustal Omega. Logoplots generated for these alignments by WebLog. In each logplot, the height of a letter is proportional to the information content of an amino acid in bits, which was calculated by equation (1) as follows.

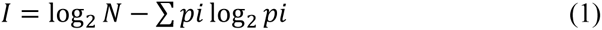

where *N* was the number of all sequences, and *pi* is the probability of the amino acid in all sequences. A large value of the unit bits indicates a high conservation of a particular site.

### Statistics

Statistical analyses were performed using GraphPad Prism 6 (GraphPad Software). EC50 values for compounds were obtained from concentration-response curves by nonlinear regression analysis. Comparation of two compounds or two constructs was analyzed by unpaired t test to determine statistical difference. All statistical data are given as mean ± SEM of at least three independent experiments performed duplicate or triplicate.

## Acknowledgments

This work was partially supported by Shanghai Municipal Science and Technology Major Project (to X.C., W.Q. and L.W.), Lingang Laboratory grant (LG202102-01-01 to X.C., LG-QS-202206-08 to L.W.), and National Natural Science Foundation of China (22007092 to L.W.).

## Author Contributions

Lifen Zhao, Methodology, Validation, Investigation, Writing – original draft preparation; Fangyu Wei, Methodology; Xinheng He, Methodology, Visualization; Hualiang Jiang, Resources, Supervision, Writing – review & editing: Liuqing Wen, Conceptualization, Methodology, Supervision, Writing – review & editing; Xi Cheng, Conceptualization, Funding acquisition, Investigation, Methodology, Project administration, Writing – review & editing.

## Supplementary files

- Supplementary file 1. Expression of mutants in HEK293.
- Supplementary file 2. Computational characterization of conformational changes and pairwise interactions of simulation models.
- Supplementary file 3. Species repertoire information for receptors.

## Data availability

All data generated or analyzed during this study are included in the manuscript and supporting file.

## Figures and figure supplements

**Figure 1—figure supplement 1.**
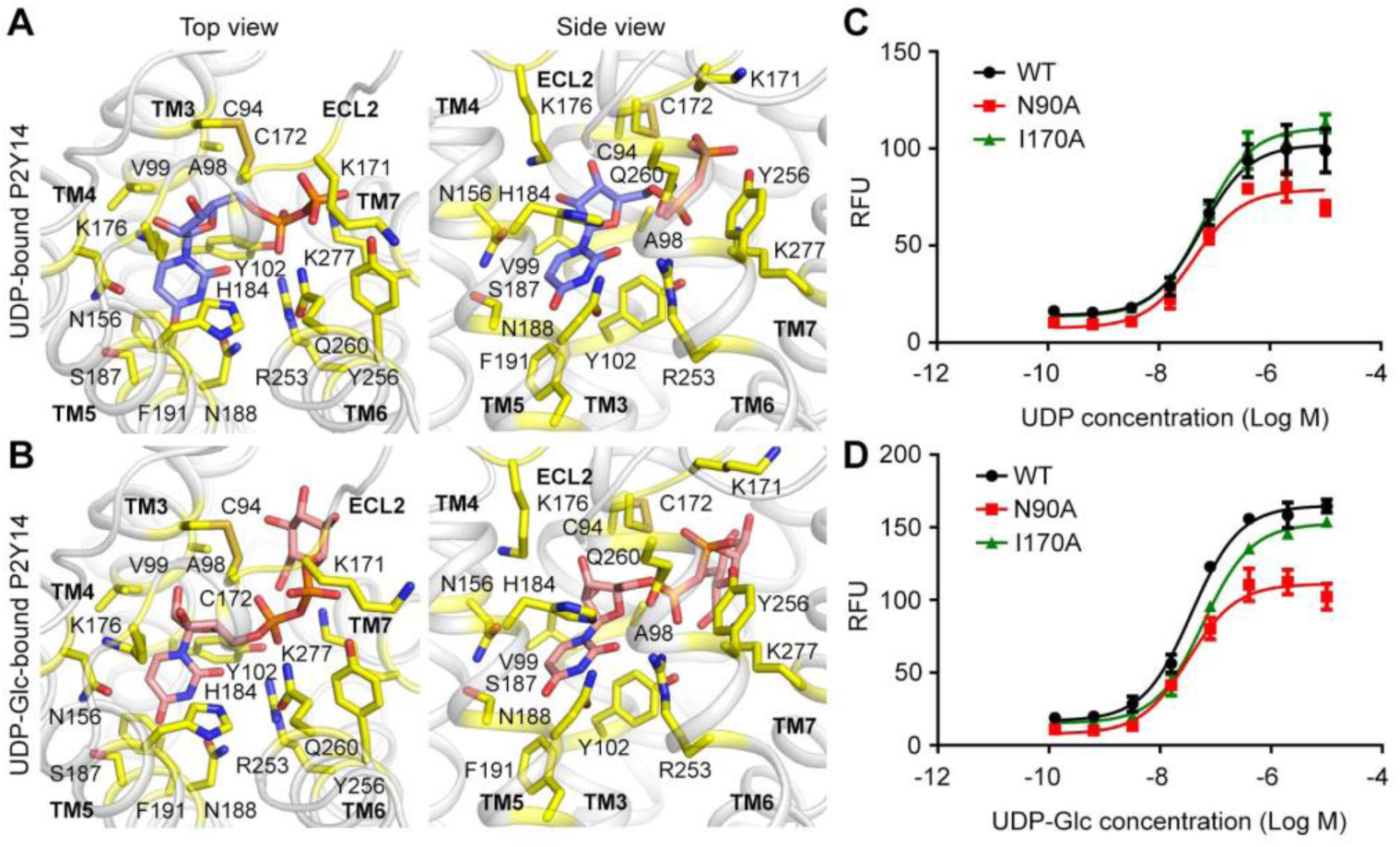
The **s**ub-pocket 1 of P2Y14 for UDP and UDP-Glc. (**A, B**) Docking models of UDP (**A**) and UDP-Glc (**B**) to P2Y14 showing binding sites for UDP moieties. Protein is shown in cartoon representation. UDP (*blue*), UDP-Glc (*salmon*) and key residues (*yellow*) are shown in stick representation. (**C, D**) Calcium mobilization concentration-response curves for UDP (**C**) or UDP-glucose (**D**) in HEK293 expressing P2Y14 WT and mutants. Data are shown as mean ± SEM (*n* = 3). See also Figure 1***—source data 2***.

**Figure 2—figure supplement 1.**
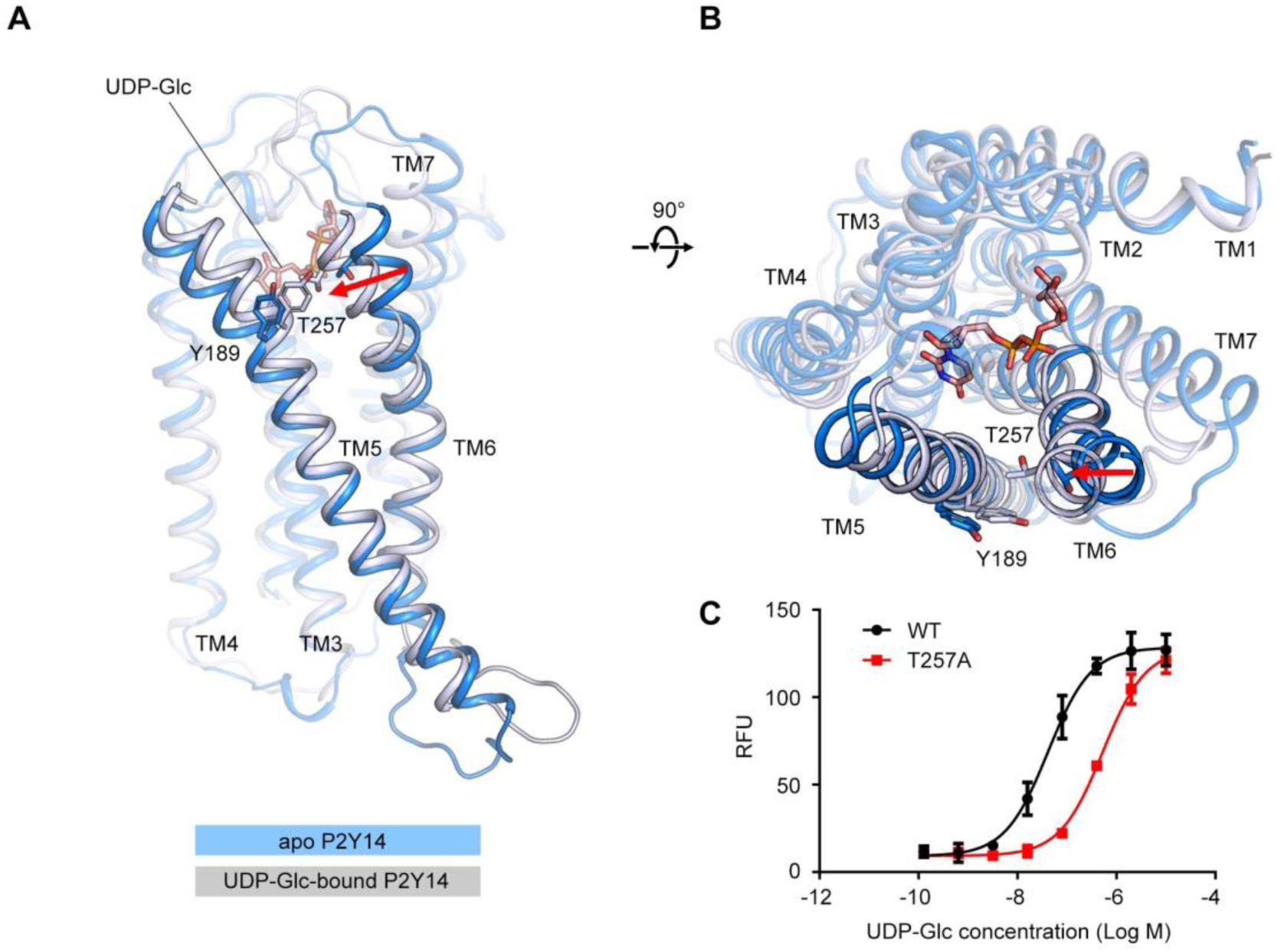
TM6 orientation in apo P2Y14 and UDP-Glc-bound P2Y14 simulation systems. (**A-B**) Side (**A**) and top (**B**) views of representative models of apo P2Y14 and UDP-Glc-bound P2Y14. Receptor is shown as cartoon. Ligand and key residues are shown as sticks. Movement of the extracellular tip of TM6 towards TM5 is shown by arrows. See ***Supplementary file 2*** for pairwise interaction details. (**C**) Concentration-response curves of calcium mobilization for UDP-Glc in HEK293 expressing P2Y14 WT and T257A mutant. Data are shown as mean ± SEM (*n* = 3). See also Figure 2***—source data 1*** and ***Supplementary file 1***.

**Figure 3—figure supplement 1.**
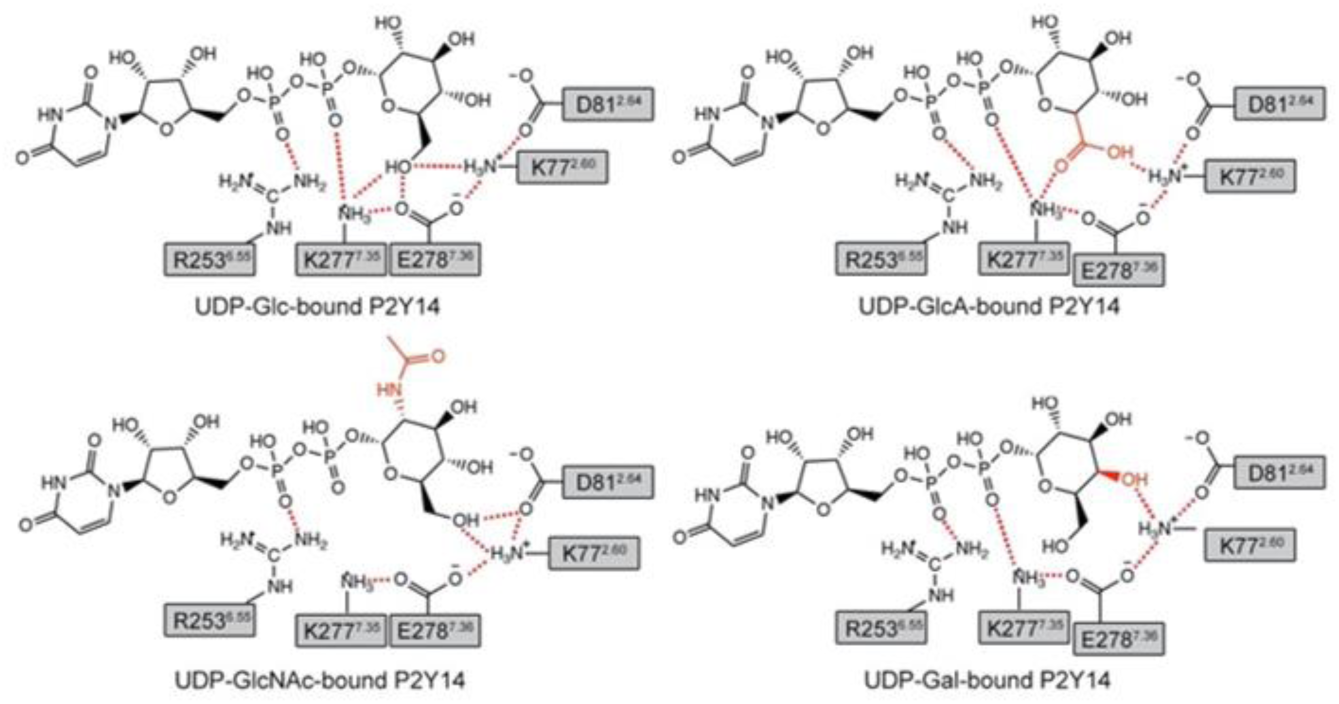
Key interactions between P2Y14 and a UDP-sugar in simulations. Putative hydrogen bonds and salt bridges are displayed as red dashed lines.

**Figure 3—figure supplement 2.**
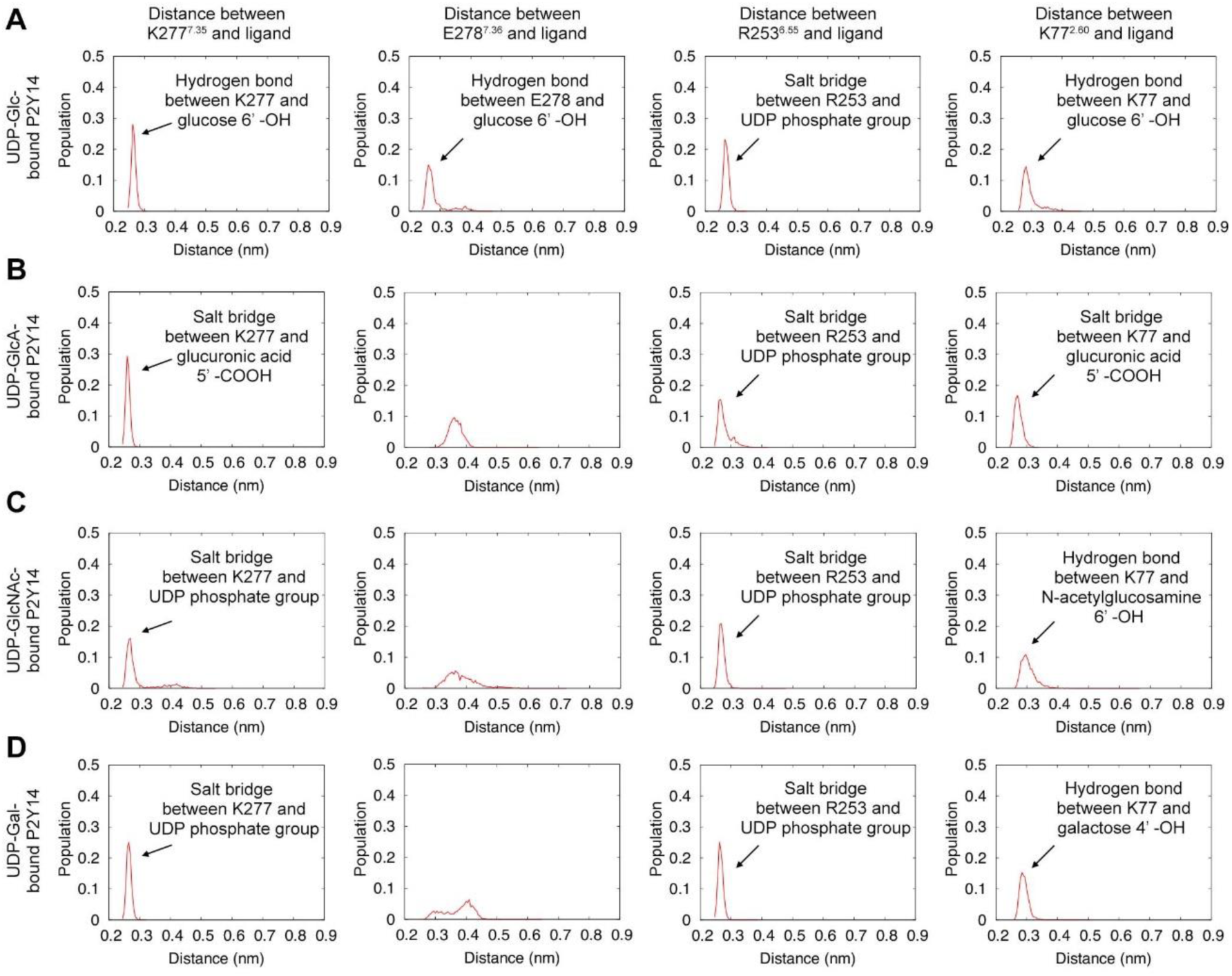
Interactions between a key residue of P2Y14 and a UDP-sugar in simulations. (**A-D**) Distribution of distance between key residues (K277, E278, R253 and K77) and a group of UDP-Glc (**A**), UDP-GlcA (**B**), UDP-GlcNAc (**C**) or UDP-Gal (**D**) in simulations.

**Figure 4—figure supplement 1.**
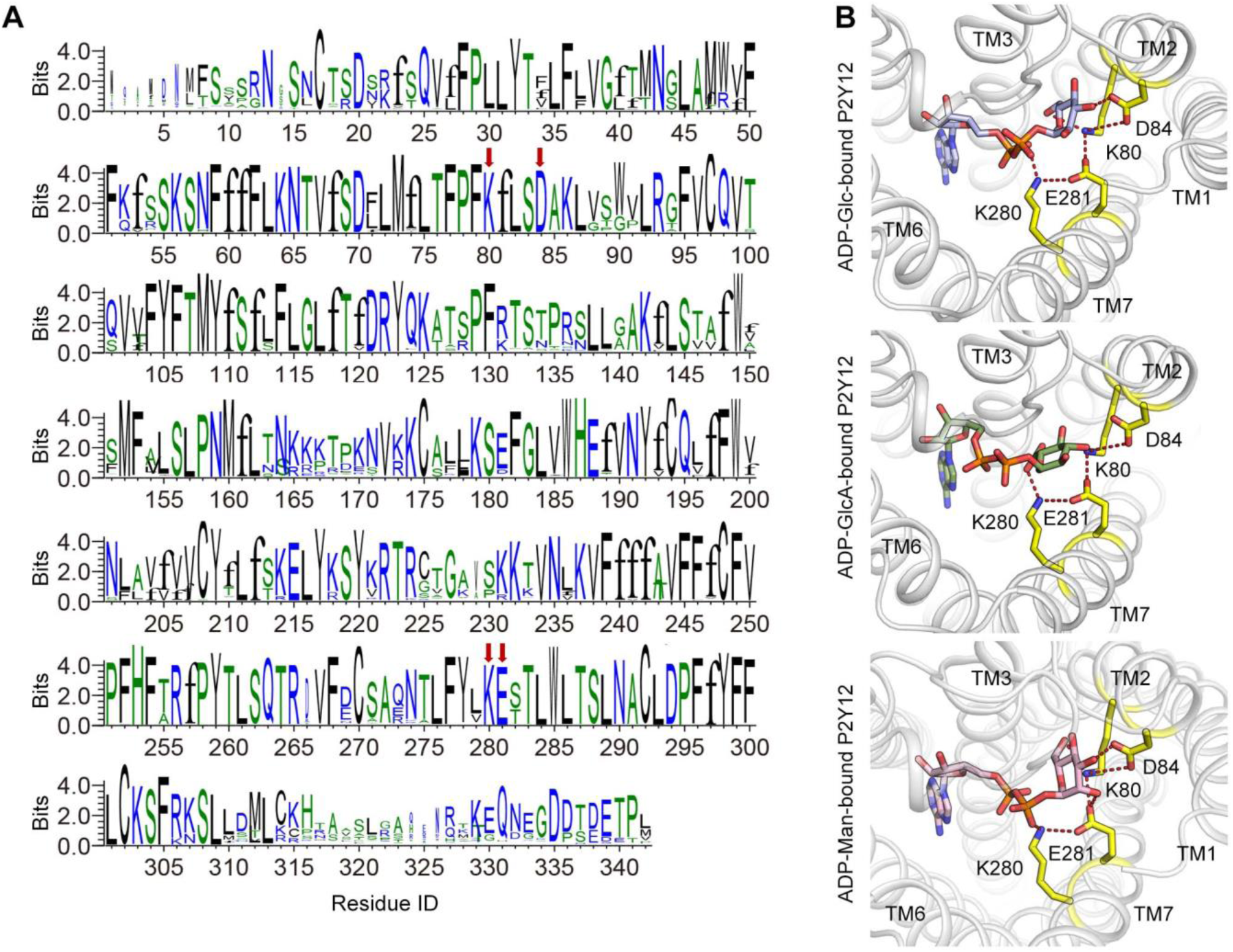
Conserved sugar-binding site on P2Y12. (**A**) Conservation of each residue on P2Y12. The height of a letter is proportional to the relative frequency of that residue at a particular site. Residues of KDKE sugar-binding motif are labeled with red arrows. See ***Supplementary file 3*** for species repertoire information. (**B**) Docking models of ADP-sugars to P2Y12. Receptor is shown as cartoon. Ligands and key residues are shown as sticks. Residues of KDKE sugar-binding motif are highlighted in yellow. Putative salt bridges are showed as red dash lines.

**Figure 4—figure supplement 2.**
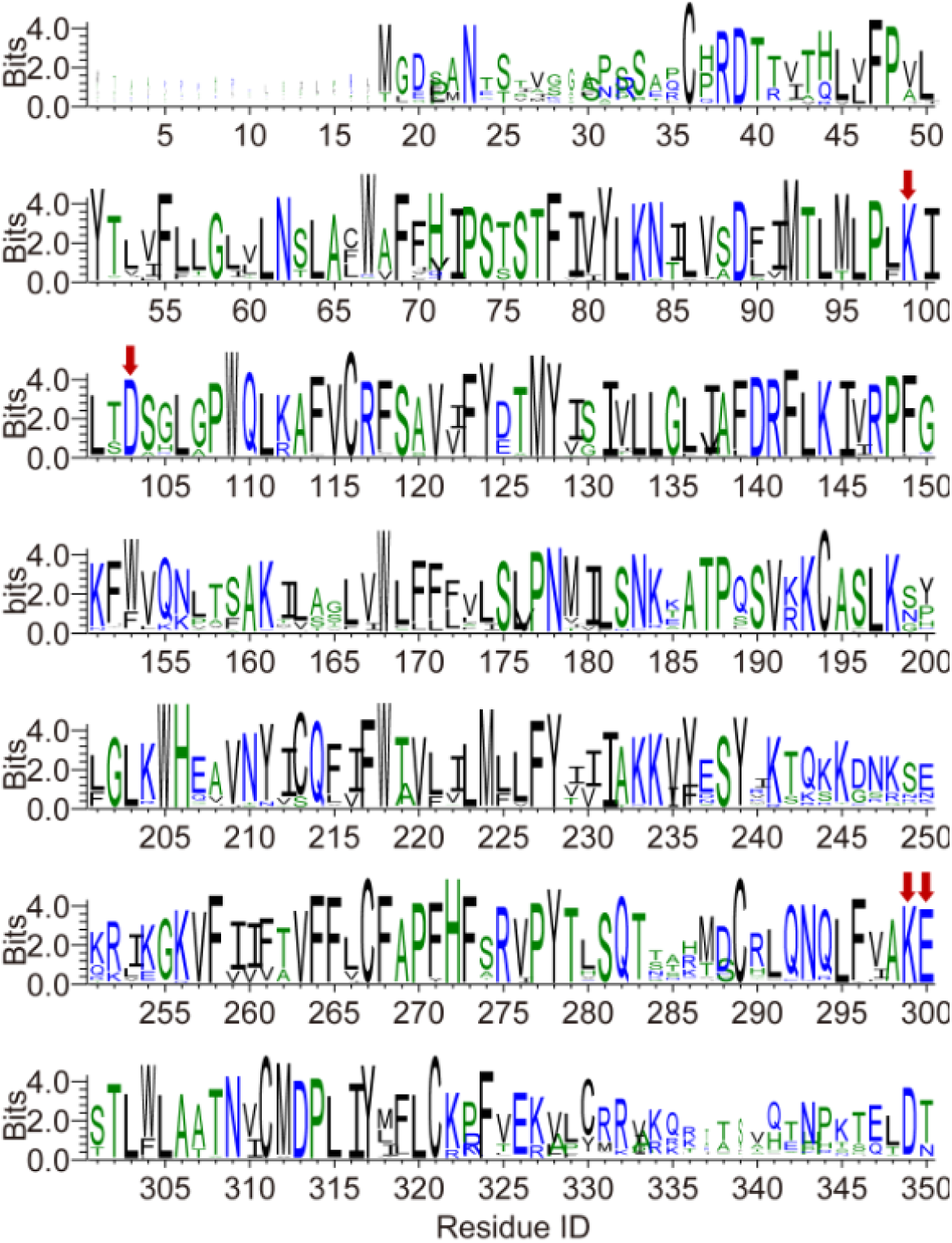
Conservation of each residue on P2Y13. The height of a letter is proportional to the relative frequency of that residue at a particular site. Residues of K^2.60^, D^2.64^, K^7.35^ and E^7.36^ sites are labeled with red arrows. See ***Supplementary file 3*** for species repertoire information.

**Figure 4—figure supplement 3.**
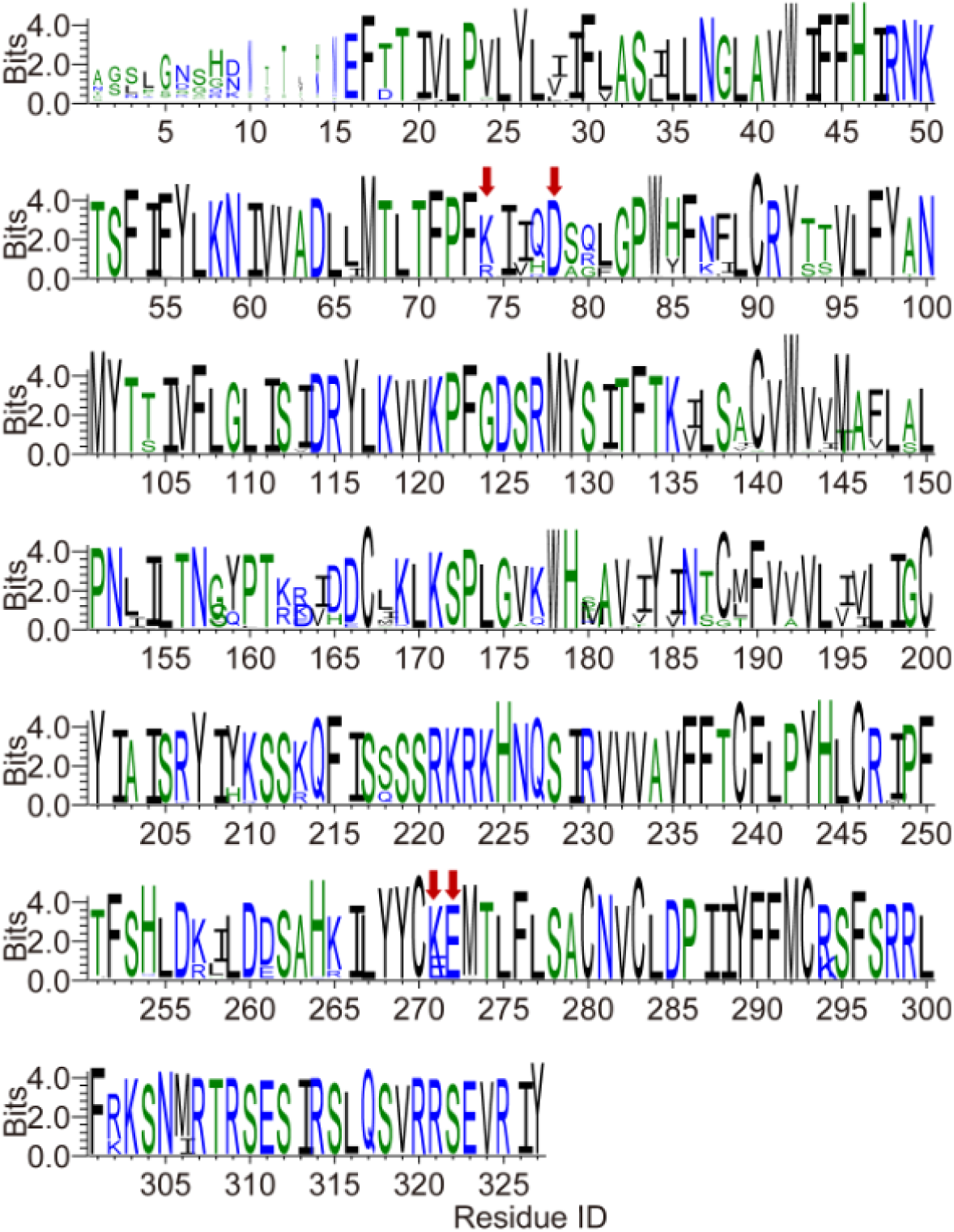
Conservation of each residue on GPR87. The height of a letter is proportional to the relative frequency of that residue at a particular site. Residues of K/R^2.60^, D^2.64^, K/E^7.35^ and E^7.36^ sites are labeled with red arrows. See ***Supplementary file 3*** for species repertoire information.

